# A general framework for cycles in ecology

**DOI:** 10.1101/2025.01.13.632680

**Authors:** Violeta Calleja-Solanas, Rafael O. Moura, José A. Langa, José R. Portillo, Fernando Soler-Toscano, Oscar Godoy

**Affiliations:** Estación Biológica de Doñana (EBD-CSIC), Spain; Instituto de Ciências Matemáticas e de Computação Universidade de São Paulo, Brazil; Dpt. de Ecuaciones Diferenciales y Análisis Numérico, Universidad de Sevilla, Spain; Dpt. de Matemática Aplicada 1, Universidad de Sevilla, Spain; Instituto Universitario de Investigación de Matemáticas (IMUS), Universidad de Sevilla, Spain; Dpt. de Filosofía, Lógica y Filosofía de la Ciencia, Universidad de Sevilla, Spain

**Keywords:** assembly graphs, asymmetry ratio, biotic interactions, ecological networks, intransitivity, species coexistence

## Abstract

Theory predicts that indirect interactions in ecological networks sustain species diversity through oscillatory dynamics. However, a framework linking interaction structure to these cycles is lacking. We develop an analytical toolbox combining assembly graphs with a mathematical decomposition of interaction matrices into symmetric and anti-symmetric components. We find that assembly cycles—closed loops of species invasions—are suppressed when symmetric interactions dominate, reflecting strong intraspecific competition. Conversely, anti-symmetric dominance, indicating competitive asymmetries, leads to cycles like rock-paper-scissors and novel multispecies invasion patterns. As asymmetries increase, new cycles emerge involving both sequential and simultaneous invasions. Applying this approach to 21 plant community interaction matrices, we find few cycles due to prevalent self-limiting effects. Our work clarifies when indirect interactions drive cycles and introduces a simple ratio assessing symmetric versus anti-symmetric contributions, constraining cycle emergence, and species coexistence in nature.

**Significant statement:** Uncertainties of the fundamental blocks that maintain biodiversity hinder our understanding of the dynamics we can observe in ecological communities. By applying a well-known mathematical approach to decomposing interactions among species within ecological communities, we discover great potential for observing rock-paper-scissors and more complex cyclic dynamics in nature. However, we predict and confirm analyzing multiple datasets that cyclic dynamics are rare because of two phenomena that pervade ecological communities. These are differences in species’ performance and the dominance of self-limiting effects. Our framework underscores the importance of studying simple properties of biotic interactions to predict complex ecological dynamics.

## Introduction

Ecologists continue to debate whether the vast biodiversity we observe in nature is maintained through mechanisms that operate between pairs of species, such as differences in phenology, natural enemies, and environmental preferences, or whether additional contributions from indirect effects within ecological networks are necessary (Soliveres, Maestre, et al. 2015; Levine, Bascompte, et al. 2017; Pajares-Murgó et al. 2024; Godoy, Stouffer, et al. 2017; Gallien, Zimmermann, et al. 2017; Bimler et al. 2024; Ranjan et al. 2024). Understanding these different mechanisms is critical for predicting the assembly and dynamics of species within communities (Wootton 1994). At one extreme, pairwise mechanisms promote a hierarchical structure of community assembly where species can coexist at a fixed equilibrium (Chesson 2000; Hofbauer and Schreiber 2022; Godoy, Soler-Toscano, et al. 2024) (Figure 1a). At the other extreme, chains of indirect interactions break such hierarchy and can lead to the emergence of cycles (May et al. 1975; Serván et al. 2021) (Figure 1b).

**Figure 1.**
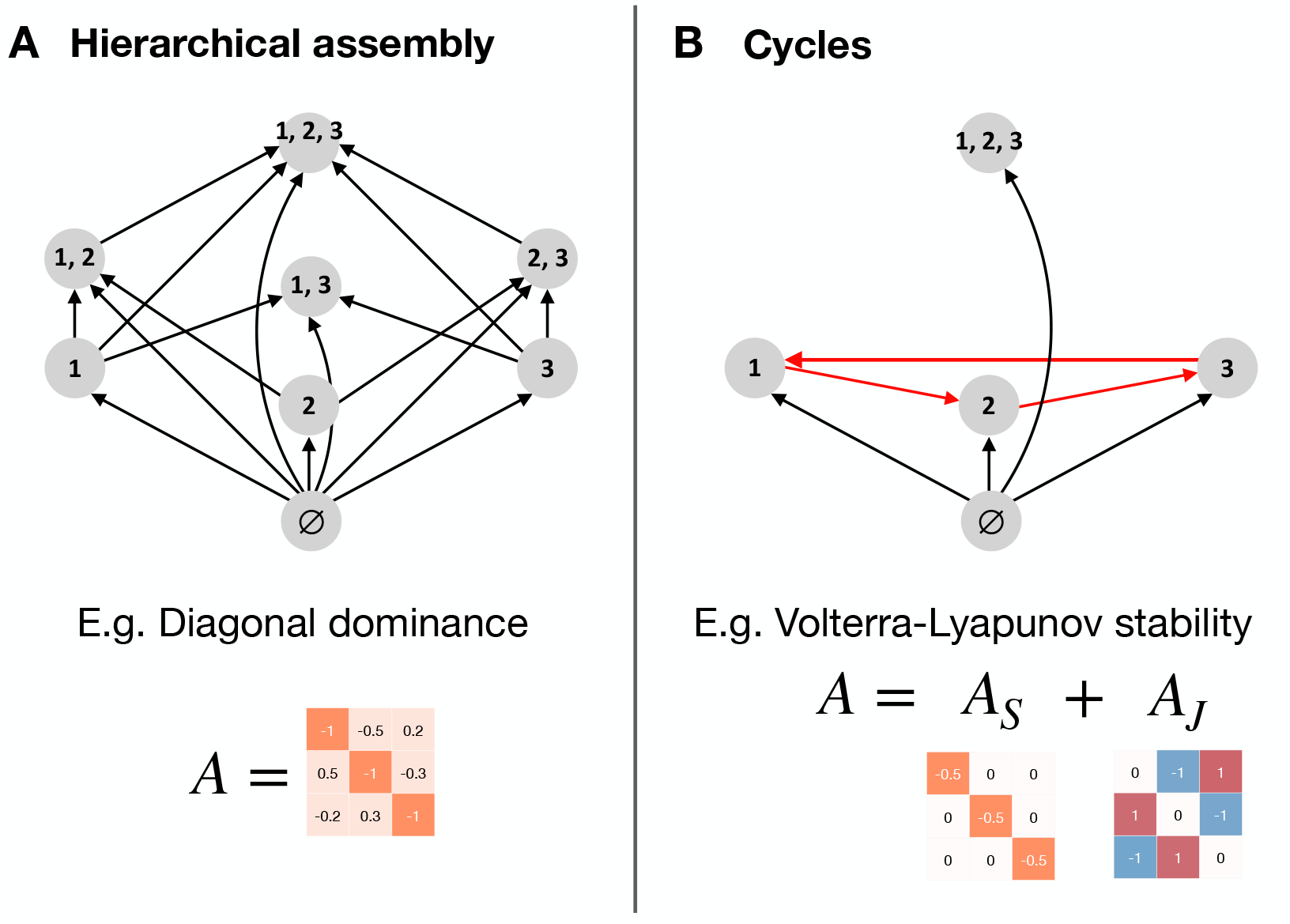
Examples of assembly graphs of 3-species systems. Nodes’ numbers represent species identity within a sub-community, and links represent possible transitions between them. The empty state with no species is denoted by ∅.(A) A hierarchical assembly structure, where a gradual invasion of one species leads to a final state. In this case, sub-community {1, 2, 3} emerges from a diagonally dominant interaction matrix. (B) Asymmetries in interaction strengths among species under Volterra-Lyapunov stability may promote cycles in the assembly graph as in the classic rock-paper-scissors game (May et al. 1975). We predict that the realization of these contrasted assembly graphs depends on the relative importance of the symmetric component of the matrix (*A*_*S*_) versus the anti-symmetric component (*A*_*J*_). Note that the structure of the assembly graph also depends on the set of intrinsic growth rates, which we will study in Figure 5.

Although this debate arises primarily from differences in environmental settings, taxa, and methodologies, there is broad consensus on the importance of the structure of species interactions (i.e., the interaction matrix) for modulating contrasted ecological dynamics (Soliveres, Maestre, et al. 2015; Gallien, Zimmermann, et al. 2017; Soliveres and Allan 2018; Hofbauer and So 1994; McCann et al. 1998; Tylianakis et al. 2010; Godoy, Stouffer, et al. 2017; Bimler et al. 2024; Ranjan et al. 2024; Song 2025). However, we still poorly understand how to map the strength and signs of species interactions onto the cycles that can occur in nature. This lack of knowledge mostly arises because the oscillatory changes in species abundances can have different origins, such as heteroclinic cycles (May et al. 1975; Krupa and Melbourne 1995; Krupa 1997), limit cycles (Gilpin 1975; Zeeman 1993; Hofbauer and So 1994), and assembly cycles (cycles arising via invasions or migration and local extinction of species) (Hofbauer and Schreiber 2022; Song 2025; Vandermeer 2025), and these kind of cycles have been have been studied separately. Moreover, the study of cycles has been restricted to the most basic configuration where no universal superior competitor exists in the interaction matrix. This configuration can lead, among other dynamics (see May et al. 1975), to the well-studied case of the rock-paper-scissors where one species becomes nearly dominant, then declines as another invades, creating oscillations in abundances (Allesina et al. 2011; Laird et al. 2006; Rojas-Echenique et al. 2011; Vandermeer 2013; Benincà et al. 2015; Alcántara et al. 2017; Matías et al. 2018; Calleja-Solanas et al. 2022). Besides intransitivity, we do not know which structure of species interactions can lead to other more complex cycles, such as multispecies invasions (Vandermeer 2025; Song 2025).

To address these limitations, we use an approach borrowed from mathematics, yet to be exploited in ecology. That is, decomposing the interaction matrix into its symmetric and anti-symmetric components (Meyer 2023; Bortolan et al. 2025), which will allow us to compare the relative importance of intraspecific and interspecific interactions. The symmetric component captures the baseline strength of interactions within the system. It captures whether community assembly is modulated by self-limiting effects arising from resource requirements or by a competitive hierarchy excluding inferior competitors. On the other hand, the anti-symmetric component captures the asymmetries in interaction strength among species, which may lead to the absence of hierarchy and then of a universal best competitor. Because the symmetric and anti-symmetric components of the matrix align well with existing ecological knowledge, they allow making clear predictions about how they contribute to the emergence of cycles. For instance, in ecological communities with strong self-limiting effects (Pajares-Murgó et al. 2024; Matías et al. 2018; Ulrich et al. 2018; Gallien, Cavaliere, et al. 2024), we predict that the likelihood of cycles is low. This is because the symmetric component of the interaction matrix dominates, while the anti-symmetric component remains negligible. Meanwhile, in ecological communities with weak self-limiting effects, and therefore, low importance of the symmetric component of the interaction matrix, we predict that the emergence of cycles depends on whether species exhibit asymmetries in their interaction strength (i.e., no competitive hierarchy). If species present asymmetries in interaction strengths (weak versus strong) and signs (positive versus negative), the presence of cycles is widespread. Some examples include marine food webs, insect communities, or plant communities in stressful and productive environments (Neutel et al. 2002; Soliveres, Maestre, et al. 2015; Benincà et al. 2015; Saiz et al. 2019; Vandermeer and Perfecto 2023; Bimler et al. 2024).

When studying cycles, it is crucial to recognize that, alongside the interaction matrix, differences in intrinsic growth rates between species are also key to defining the dynamics of ecological systems. The inherent ability of species to grow significantly influences their ability to recover from low density, withstand environmental variability, and ultimately, the coexistence of entire communities (Chesson 2000; Gallien, Zimmermann, et al. 2017). Most past theoretical work has studied cycles when intrinsic growth rates are alike between species (May et al. 1975; Laird et al. 2006) for simplicity reasons. Recent theoretical work and empirical evidence suggest that cycles are more likely to occur when the variability in species’ intrinsic growth rates is lower (Soliveres, Maestre, et al. 2015; Godoy, Stouffer, et al. 2017; Gallien, Landi, et al. 2018; Matías et al. 2018; Saiz et al. 2019; Ranjan et al. 2024), that is when the innate ability of species to produce offspring and survive is similar among species. Together, these previous findings posit that the likelihood of cycles decreases as differences in species’ intrinsic growth rates increase. However, an exploration of this prediction in combination with the interaction matrix is currently lacking.

Here, we introduce a framework capable of predicting the presence of assembly cycles of any length and number of species between transitions. We believe this framework is paramount to enriching our understanding of the prevalence of cycles in nature and predicting assembly structures that have not yet been discovered. We focus on assembly cycles since they are the only ones that show a direct correspondence between a cycle in the interaction matrix, a cycle in the assembly graph, and cyclic dynamics in the temporal behavior of the system. Additionally, focusing on the assembly of communities through assembly graphs has the benefit that incorporates the combined effects of species’ intrinsic growth rates and interactions matrix on community dynamics (Serván et al. 2021; Hofbauer and Schreiber 2022; Almaraz et al. 2024). Briefly, an assembly graph departs from an empty community and indicates the different transitions among sub-communities (i.e., subsets of species that coexist, a stationary solution of our model) until the community reaches a final non-invasible community. These sub-communities, represented by nodes, are depicted at different levels according to their number of species (Figure 1).

By examining the conditions under which cycles arise in this graph —considering the ratio of symmetry to asymmetry in interactions and the variation in intrinsic growth rates— we discover three main groups of cyclic structures: (1) *Level-1 cycles* or cycles of sub-communities of a single species, that is, generalizations of the rock-paper-scissors dynamics for any number of species (Figure 1a). (2) *Level-ℓ cycles* cycles, which are composed of sub-communities of *ℓ* species and allow for simultaneous invasions of more than one species. (3) *Multi-level cycles* in which the sub-communities are composed of a varying number of species. Beyond these three categories, we found that cycles are more likely to occur when the anti-symmetric component dominates the interaction matrix and species show similar intrinsic growth rates, regardless of community richness. To confirm that this combination is rare in nature, we apply our framework to a compilation of existing datasets of interaction matrices in the literature that covers a wide range of species richness and plant communities across contrasted climatic conditions (see Adler et al. 2018 for details). As expected, we found low evidence of cycles in such empirical data. This last result opens the discussion of the ecological constraints, both internal (i.e., species interactions) and external (i.e., environmental conditions), on cycles in natural systems.

## Methods

The core of our analysis uses assembly graphs to study cycles in different structures of community assembly (Serván et al. 2021; Hofbauer and Schreiber 2022). The nodes of an assembly graph are subsets of coexisting species, technically known as sub-communities or admissible communities. Nodes are connected by directed links when there is a potential transition due to invasions by missing species (Figure 1).

Within the framework of assembly graphs, a cycle is defined as a closed loop that represents sequences of species invasions that eventually return to the initial sub-community, generating oscillations in species abundances. This sequence of invasions is encoded directly in the topology of the assembly graph (red lines in Figure 1b) and reflects temporal turnover driven by species arrivals. Specifying this definition is important as the term “cycle” refers to different concepts in the ecological literature, particularly in studies of intransitivity and community assembly. Periodic orbits in the phase space, such as limit cycles, are also plainly referred to as cycles. They also generate oscillatory population dynamics, but they are not driven by sequential invasions. They rather emerge from intrinsic population dynamics within a sub-community — for instance, due to predator-prey interactions. Then, these dynamical cycles can also be contained in general within a node of the assembly graph, but they are not represented as a cyclic graph. They “live” inside a single individual node of the assembly graph, and their detection can only be revealed by performing either direct simulations of population dynamics or mathematical analysis of the sub-community’s stability properties (Takeuchi 1996; Bunin 2017). In sum, assembly cycles are the only ones that show a direct correspondence between a cycle in the interaction matrix, a cycle in the assembly graph, and cyclic dynamics in the temporal behavior of the system.

We model the population of *n* species through the generalized Lotka-Volterra equations, where each species interacts positively and/or negatively. This set of nonlinear differential equations describes the changes in the density *x*_*i*_ of species *i* over time as:

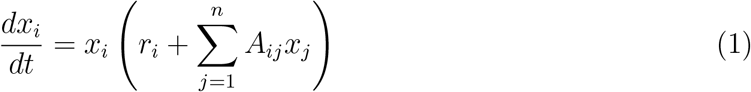

The vector or intrinsic growth rates *r* = (*r*_*i*_) denotes species performance, i.e. how species would grow isolated under given abiotic conditions. In turn, interactions among species are phenomenologically captured by the *n* × *n* interaction matrix *A*, where *n* denotes the total number of species. Its diagonal elements *A*_*ii*_ represent intraspecific interactions, that is, how a species limits itself. Interspecific competition is captured in *A*_*ij*_, the effect of species *j* on the per-capita growth rate of species *i*. The interaction matrix can be expressed as *A* = *A*_*S*_ + *A*_*J*_, the sum of a symmetric *A*_*S*_ and anti-symmetric *A*_*J*_ components (Figure 1b), where 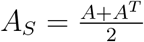 and 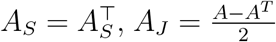 and 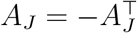. To analyze how an interaction matrix *A* determines an assembly of cycles, compute the asymmetry ratio:

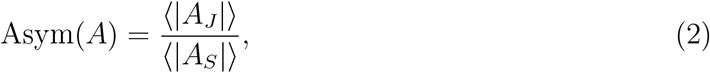

defined as the ratio between the magnitude (strength) of the entries of the anti-symmetric ⟨|*A*_*J*_ |⟩

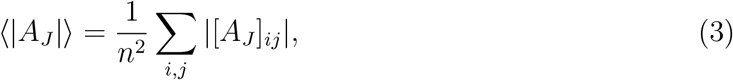

and symmetric

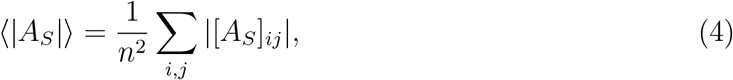

components of the interaction matrix. Such decomposition covers a wide range of ecological phenomena: when intraspecific competition dominates, i.e., large ⟨|*A*_*S*_|⟩ as in diagonally dominant matrices, Eq. (6), communities stabilize in the absence of cycles. On the other hand, systems with greater asymmetry of interactions, i.e., larger ⟨|*A*_*J*_ |⟩, create directional imbalances that can promote complex cyclic behavior.

Since the connections between the nodes of an assembly graph are established via successful invasions, in this model, the invasion rate of the species *i* on the sub-community *I* is the average species per-capita growth rate when rare, defined as:

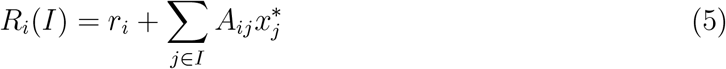

If the sub-community is empty, *I* = ∅, we use the convention *R*_*i*_(∅) = *r*_*i*_ (Hofbauer and Schreiber 2022). The graph contains a link from a sub-community *I* to a sub-community *I*^′^ if, and only if, *I* ≠*I*^′^, and *R*_*j*_(*I*) *>* 0 for every species *j* that is in *I*^′^ but not in *I*, and *R*_*i*_(*I*^′^) *<* 0 for every species *i* that is in *I* but not in *I*^′^. This indicates that the conditions in *I* allow the invading species *i* to establish and persist. Conversely, for the transition to occur, any species that was in the initial sub-community *I* but not in the new one *I*^′^ cannot invade it. Because the AG is a directed network, cycles emerge when a path of directed links starts and ends at the same node. The main benefit of using assembly graphs is that they define invasions considering both the interaction matrix *A* and intrinsic growth rates *r*. Therefore, assembly graphs are a powerful tool for studying the synergies between species interactions and performance in cycle formation.

The framework is flexible enough to explicitly describe the structure of all connections between equilibria, but we restrict our study to *Volterra-Lyapunov stable* (VL) matrices. There are two reasons for this choice. The practical reason is that the links in the assembly graph are indeed connections in the phase space of the Lotka-Volterra system (Almaraz et al. 2024) for the broad spectrum of interaction matrices that meet the condition of Volterra-Lyapunov stability, Eq. S1. Secondly and critically, VL-stable matrices contain an ecologically relevant subset of matrices, *Diagonally Dominant* (DD) matrices, which satisfy:

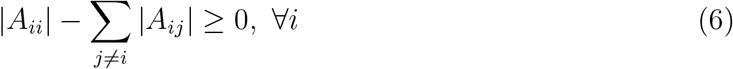

In a DD matrix, self-regulation plays a stronger role than interspecific competition, which might reflect differences in resource use among species (Tilman 1982). The DD matrices always have an asymmetry ratio of less than one by definition. Mathematically, diagonal dominance can also be defined by columns:

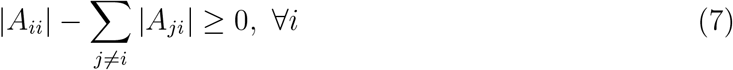

Both definitions are closely tied to competition asymmetry, as species with stronger self-limitation are less likely to exclude others, promoting coexistence and reducing the dominance of a single fast-growing competitor (Levine and HilleRisLambers 2009; Daniel et al. 2024).

To systematically explore the conditions under which assembly cycles emerge, we coupled our theoretical framework with an algorithm recently developed (Bortolan et al. 2025). This method serves two main goals. First, it generates random VL-stable matrices that allow exploring the assembly graph for any number of species (Appendix S1). Second, it also generates interaction matrices tailored to include simultaneous invasions of one or more species (Appendix S2), which expands beyond the traditional approach of considering cycles between single-species equilibria (May et al. 1975; Hofbauer and So 1994; Gallien, Zimmermann, et al. 2017; Vandermeer 2025; Song 2025).

## Results

We show that the presence of cycles in ecological systems depends on the asymmetry ratio of the interaction matrix *A*. Following our predictions, the dominance of the anti-symmetric component of interactions promotes many different types of cycles, while the preponderance of the symmetric component inhibits them (Figure 2a). Therefore, the probability of cycles increases with the magnitude of the anti-symmetric component ⟨|*A*_*J*_ |⟩ and decreases with the magnitude of the symmetric component ⟨|*A*_*S*_|⟩. At one extreme, an ecological system has a high probability of containing cycles when the symmetric component is weak compared with the anti-symmetric component, especially for systems with high richness. At the other extreme, such probability goes to zero when the anti-symmetric component tends to zero, and therefore the interaction matrix becomes symmetric. There is a special case in which this probability decreases sharply at a ratio that we have computationally found to be Asym(*A*) = 1. This particular case includes diagonally dominant matrices. This limitation of cycles is explained because the self-regulation rates *A*_*ii*_ are in the symmetric component of *A* (the diagonal of *A*_*S*_ is the diagonal of *A*, and the diagonal of *A*_*J*_ is zero). In other words, when *A* is diagonally dominant, its symmetric component is always larger than its anti-symmetric component, so diagonally dominant matrices are located before the region where the asymmetry ratio equals one. The mathematical proof describing why diagonally dominant matrices do not contain any type of cycle is currently unknown except in particular cases, for instance when cycles are among sub-communities of one single species (Bortolan et al. 2025). However, we believe our result is robust because no cycles in assembly graphs were found by sampling millions of diagonally dominant matrices coupled in a Lotka-Volterra model with positive and negative intrinsic growth rates. In conclusion, our results suggest that diagonally dominant matrices are incompatible with the presence of any kind of cycle.

**Figure 2.**
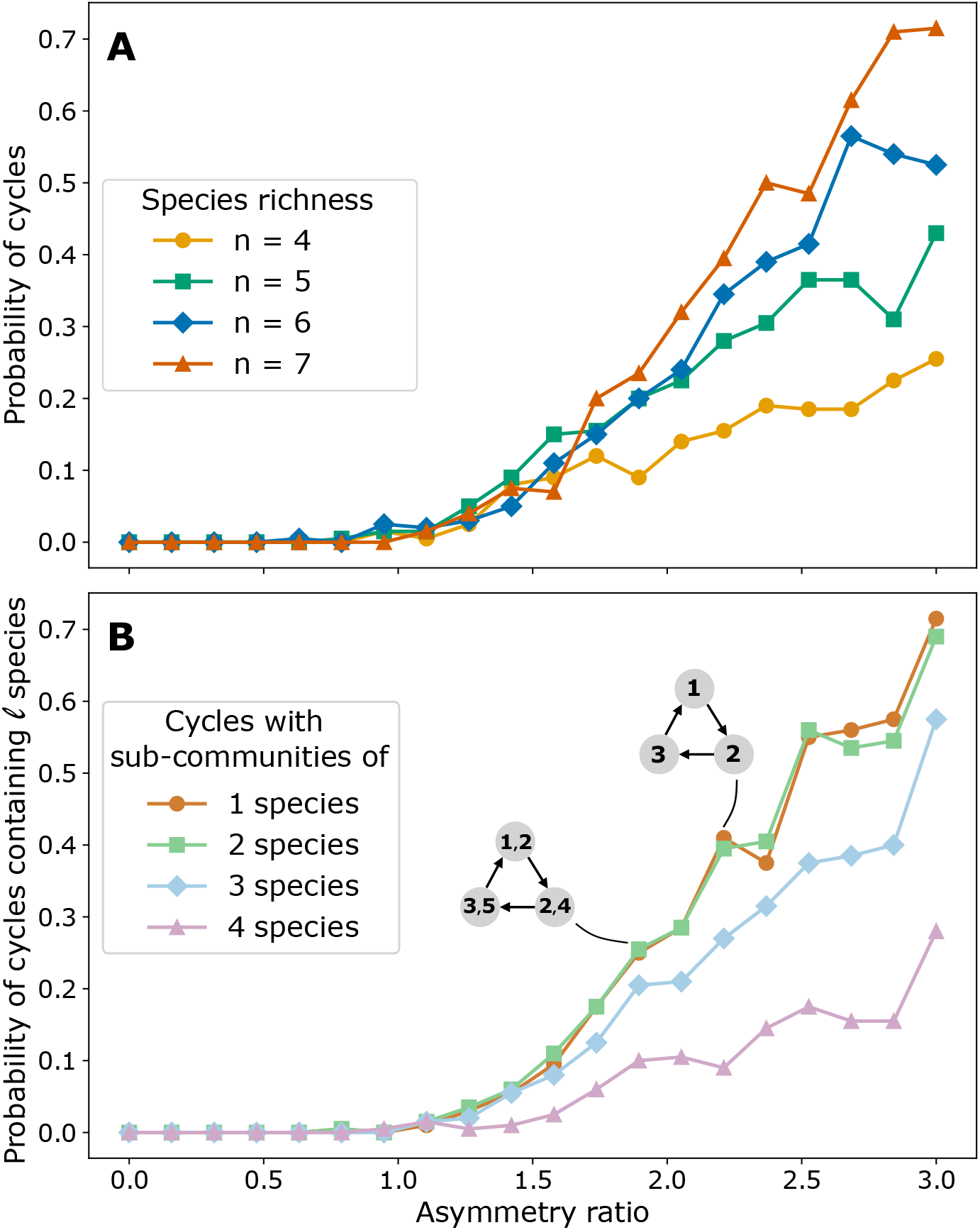
Influence of the asymmetry ratio on the probability of cycles. For 20 values of the asymmetry ratio, defined in Eq. (2), with a range in species richness, we sampled 200 matrices *A* randomly and calculated the associated assembly graphs for a total of 16.000 matrices. (A) The probability of an assembly graph containing any type of cycle increases non-linearly with the asymmetry ratio and with the species richness. That is, the greater the asymmetry ratio, the more likely it is to find a cycle; and, for a fixed asymmetry ratio, the larger the potential number of species, the more likely to find a cycle. (B) The probability of cycles whose nodes contain a fixed number of species also increases with the asymmetry ratio. Overall, cycles formed by sub-communities of a single species (class A in Figure 3 and top graph) are more likely to occur than cycles containing 2, 3, or 4 species in each sub-community (class B in Figure 3 and bottom graph). Intrinsic growth rates are all equal to one for both panels.

Our results also show that when the anti-symmetric component of the matrix dominates, we obtain three distinct classes of cycles: single-species, multispecies, and species-varying cycles (Figure 3). The first category encompasses a generalization of the rock-paper-scissors to any number of species. This generalization means that cyclic dynamics following the May-Leonard case of sequential transitions (May et al. 1975) of single species between sub-communities can be described for assemblages of any odd number (Figure 3a) as previous work suggested (Allesina et al. 2011; Vandermeer 2011). Nevertheless, this generalization also encompasses cycles with an even number of species, which has been much less studied before. Overall, this first category includes any assemblage of species that contains an assembly cycle of sequential invasions of an arbitrary number of species length. Such cycles can contain all species within the assemblage or only a subset of them (Figure 4). The other two categories correspond to cycles with more complex transitions beyond the traditional single-species sub-communities. In more detail, the second category (*multispecies cycles*) corresponds to cycles between states with more than one species. Figure 3b shows the rock-paper-scissors-lizard-Spock system, a generalization of the rock-paper-scissors for a richness of five species, where each sub-community has three species sequentially invaded by one species at a time. Finally, the cycles in the third category (*species-varying* cycles) can involve not only sequential but also simultaneous multispecies invasions, transitioning between sub-communities of a varying number of species. Figure 3C shows an example of a cycle transitioning between sub-communities composed of either one or two species. Finally, we also found that cycles composed solely of one-species equilibria (the first category) are more common than cycles containing at least a sub-community of *m < n* species, which correspond to the latter categories (Figure 2b).

**Figure 3.**
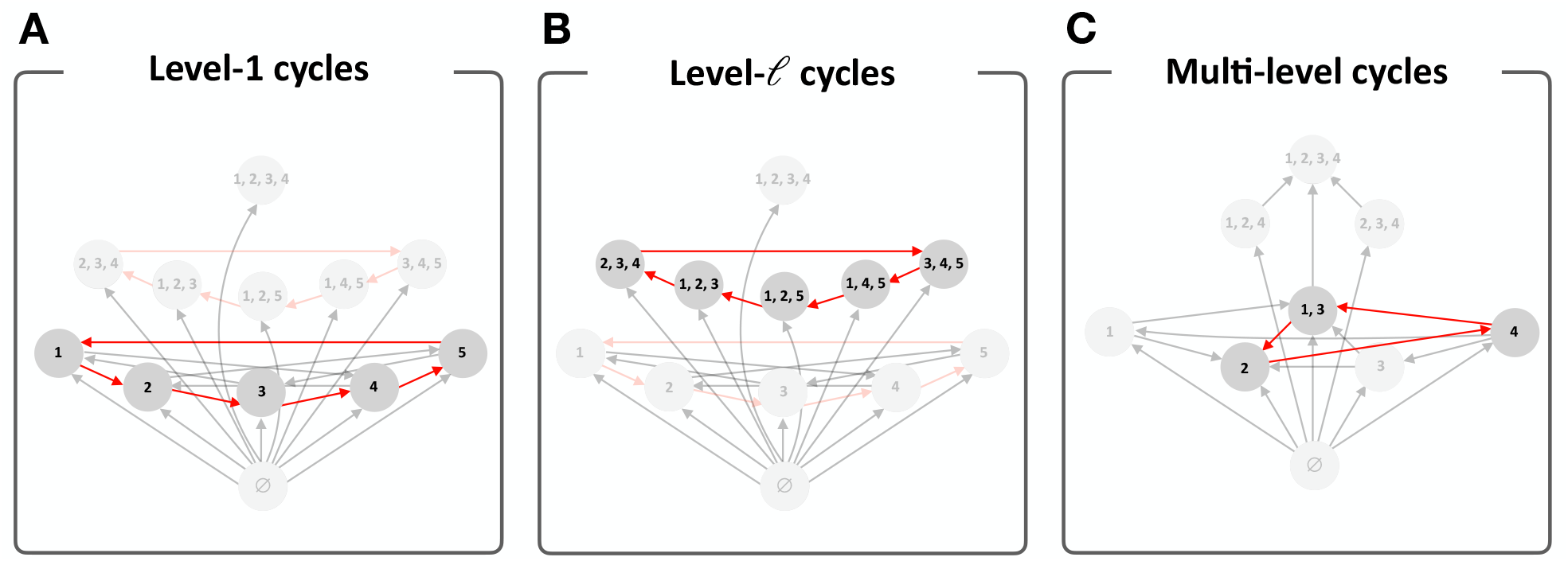
Categories of cycles: (A) Level-1 cycles: cycles among one-species sub-communities, which can be seen as generalizations of the rock-paper-scissors dynamics. Here is the assembly graph of the rock-paper-scissors-lizard-Spock system, which involves sequential invasions among five species. (B) Level-*ℓ* cycles: Cycles involving a fixed number of species *ℓ >* 1 in their sub-communities. (C) Multilevel cycles: cycles with a heterogeneous number of species among sub-communities. We illustrate here a particular case of sequential invasions of one species and a simultaneous invasion of two species (from sub-community {4} to {1, 3}). These three cycles, highlighted in red, have been built with an algorithm of cycles *à la carte* (Appendix S2).

**Figure 4.**
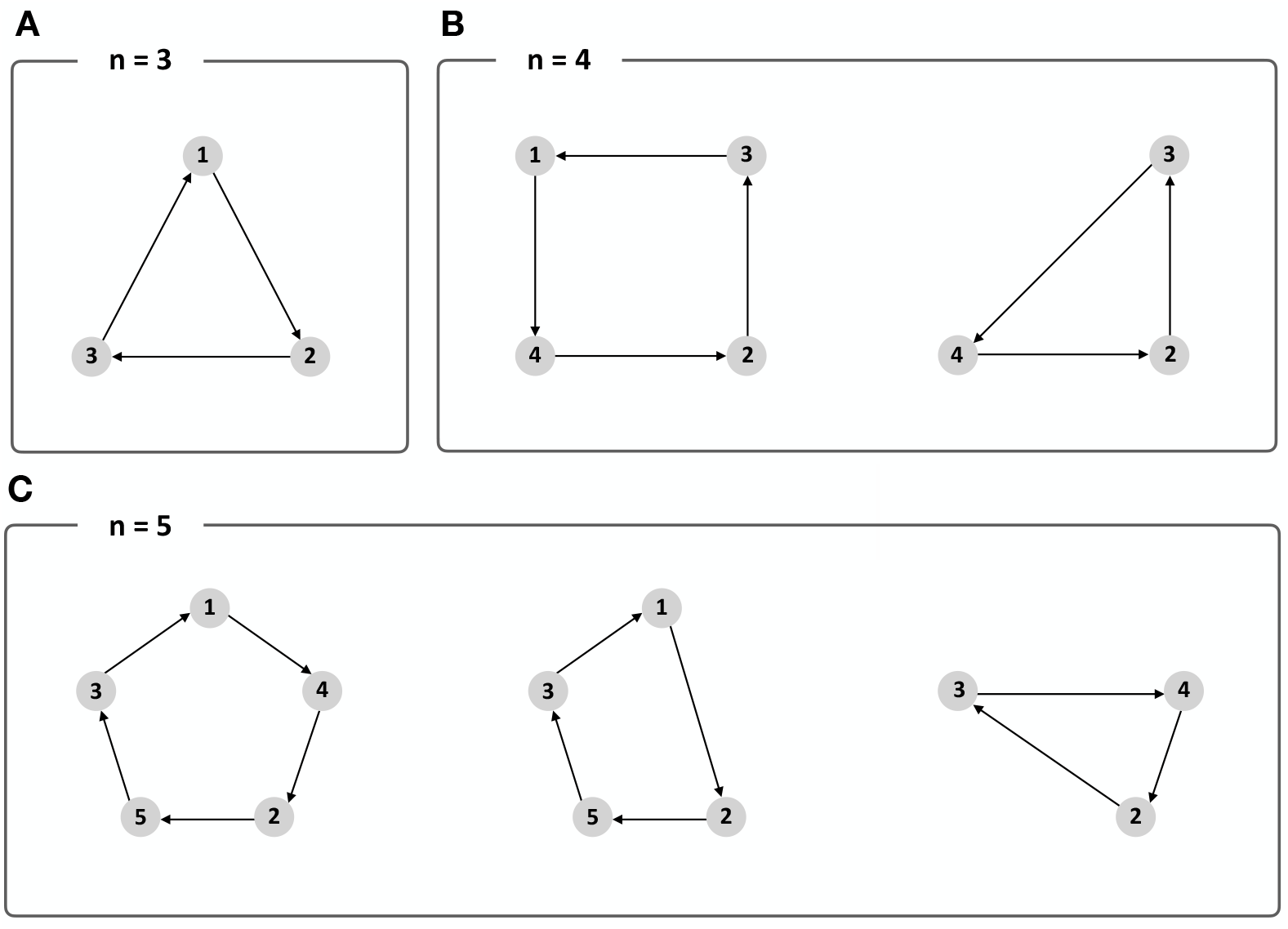
Examples of generalizations of the rock-paper-scissors system (A) to four (B) and five (C) species, where level-1 cycles of an arbitrary length can emerge. In the panels, the assembly graphs are not depicted but some of the present cycles.

To study the effect of the anti-symmetric component of the matrix in isolation, the values of the intrinsic growth rates in Figure 2 were all taken equal to one. Nevertheless, as theoretical and experimental work has shown before, the probability of finding a cycle in the assembly graph also depends on the variation of the intrinsic growth rates (*r*). For our study, and in agreement with previous findings, we found that the probability of cycles is inversely proportional to the variability of *r* (Figure 5a). This means that cycles are more frequent when variation in intrinsic growth rates is minimal. Conversely, there are fewer cycles in an assembly graph when intrinsic growth rates are more variable, including both positive and negative values. When combining the effect of both the asymmetry ratio and the variation in intrinsic growth rates, we discovered that the highest probability of finding a cycle in an assembly graph corresponds to a low variation in intrinsic growth rates and a high asymmetry ratio. If we focus now on the complexity of the assembly graph, understood as the number of sub-communities involved in the cycle, we find an opposite pattern for the asymmetry ratio. This means that more symmetric interactions produce a larger number of sub-communities at equilibrium in the assembly graph, and therefore, enlarge the number of possible invasion paths to coexistence (Figure 5b). For intrinsic growth rates, lower variation also promotes more complexity of the assembly graph. Therefore, the assembly graphs with more frondosity (i.e., transitions between sub-communities) emerge under the combination of low asymmetry ratio and low variation in intrinsic growth rates.

**Figure 5.**
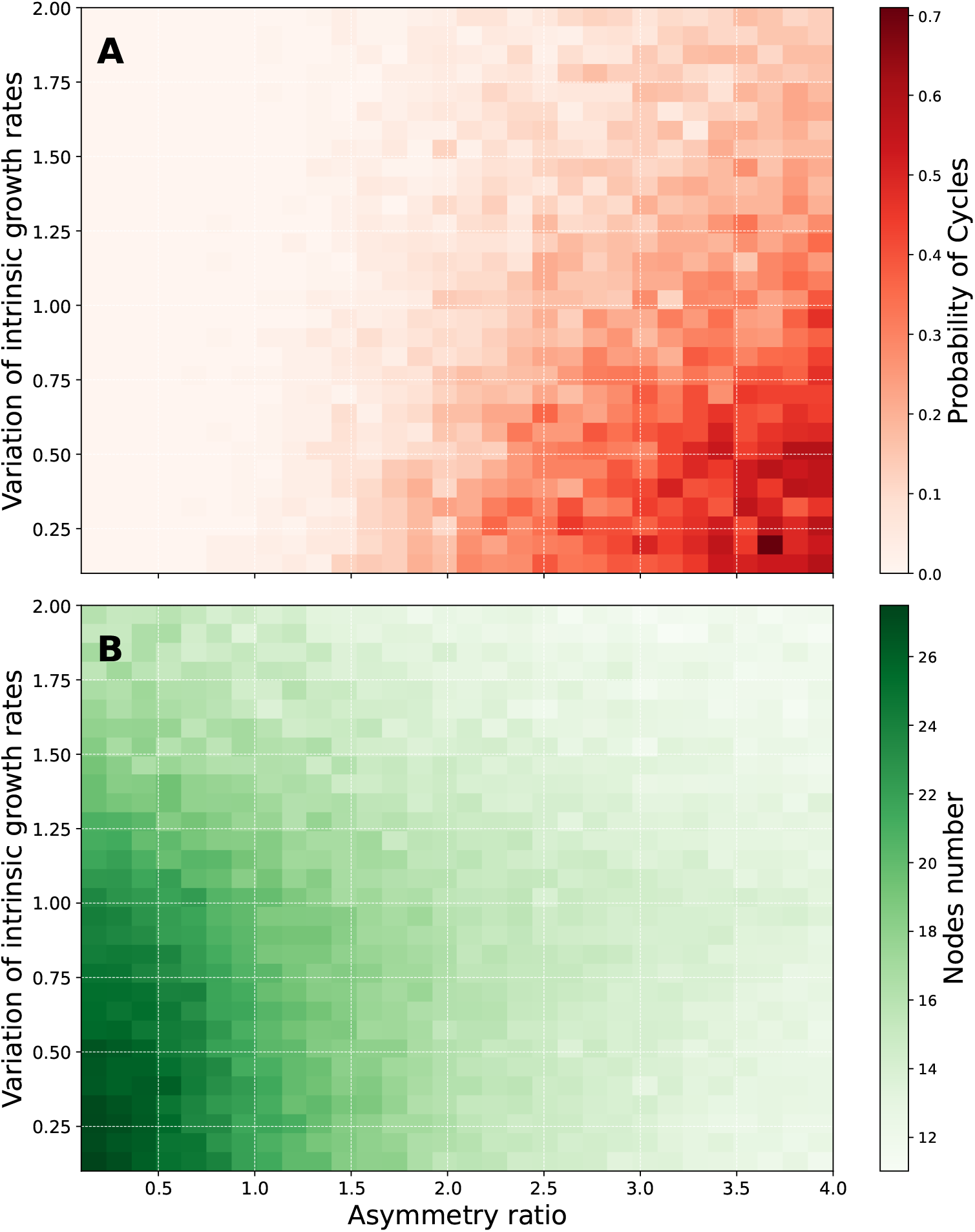
(A) The color gradient captures the probability of finding cycles in the assembly graph, given both the variation in the asymmetry ratio and species’ intrinsic growth rates. The y-axis represents the variation limit of the intrinsic growth rates, centered on the value of 1. Then, for a value of 0.4 the intrinsic growth rates are sampled uniformly at random between 0.6 and 1.4. Notice that high values of variation (larger than 1 in our case) allow negative growth rates. (B) The color gradient represents the mean number of sub-communities of the assembly graphs. The number of assembly graphs tested for each combination of parameters was 100 and had a maximum of 5 species.

Finally, to explore our general framework beyond simulations, we evaluated the potential emergence of cycles in 21 empirically determined matrices that account for a wide variety of terrestrial plant communities with varying levels of species richness (Adler et al. 2018). We found that only 8 (38%) of these matrices are diagonally dominant either in a row or column sense, that is, the strength of intraspecific interactions is stronger than interspecific interactions, Eq. (6). Moreover, a greater proportion of them (14 matrices, 66%) also meet the more strict conditions of Volterra-Lyapunov stability, Eq. S1. In contrast, the remaining third of the matrices could not be classified as VL-stable (33%). Once their stability was assessed, we tested for the presence of cycles following two complementary approaches. These approaches cope with the fact that information on the variation in intrinsic growth rates between species is missing. On the one hand, we built assembly graphs considering all intrinsic growth rates between species equal to one. We found no cycles for any community, instead, 5 of them (24%) presented full frondosity, allowing the existence of any possible sub-community. On the other hand, we sampled a thousand random vectors of intrinsic growth rates within the interval (−1, 1) for each of the empirical matrices. We found that the occurrence of cycles in assembly graphs was associated with only 2 of the 21 total communities (9%) for some values of intrinsic growth rates. This negligible proportion agrees with the fact that the asymmetry ratio across the 21 communities is low, with an average of 0.41 and a variance of 0.06.

Taken together, the combination of results obtained from simulations and their application to empirical interaction matrices indicates that there is, in principle, a strong potential for the emergence of assembly cycles in ecological systems. These cycles can involve the well-documented rock-paper-scissors, but there is also potential for much more complex assembly dynamics. However, the realization of these cycles seems unlikely in nature, at least for the dataset analyzed. This is because of the restriction imposed by the biotic interactions among species, in which the relative importance of self-limiting effects compared to interspecific effects (i.e., a tendency towards diagonally dominant matrices) reduces the anti-symmetric component of the matrix, and hence the presence of cycles.

## Discussion

Despite decades of research, ecologists still strongly debate the prevalence and mechanisms of cycles in nature (Laird et al. 2006; Soliveres and Allan 2018; Gallien, Zimmermann, et al. 2017; Godoy, Stouffer, et al. 2017; Adler et al. 2018; Bimler et al. 2024). This debate is more broadly linked to the importance of mechanisms of biodiversity maintenance that operate between pairs of species versus indirect chains of interspecific interactions (May et al. 1975; Chesson 2000; Levine, Bascompte, et al. 2017; Gallien, Zimmermann, et al. 2017). Here, we present a general framework that captures the essence of this discussion and show with an associated mathematical toolbox a metric that predicts the presence or absence of assembly cycles for any given ecological community composed of an arbitrary number of species. This metric, called asymmetry ratio, summarizes the balance between the symmetric and the anti-symmetric components of a matrix containing the interaction strengths and signs within and between species. In more detail, our framework expects the presence of cycles when the anti-symmetric component in the interaction matrix dominates, Asym(*A*) = ⟨|*A*_*J*_ |⟩*/*⟨|*A*_*S*_|⟩ *>* 1, but restricts them when the symmetric component dominates, Asym(*A*) *<* 1. In ecological terms, this means that cycles are absent when the assembly of a community is governed by stronger intraspecific than interspecific interactions. Alternatively, cycles are present when intraspecific interactions are overall weak and there are strong asymmetries in interspecific interactions. This result is a novelty that completes the study of cycles and symmetric interactions. MacArthur 1969 found a Lyapunov function that excludes cycles in symmetric systems, but it had not been proven that asymmetries interactions promoted cycles. Overall, our framework shows that the complexity of biotic interactions we observe in ecological communities can be summarized in a simple asymmetry ratio that predicts the presence of cycles in nature.

The study of cycles in ecology has always dealt with sequential changes in species dominance and hence in their abundance. Our study enlarges this current view, showing that these sequential dynamics are only a small part of the picture. With larger species pools, assembly cycles can be much more complex, including cycles with multispecies invasions or cycles with varying species invasions. Whether this occurs in nature remains to be empirically described. Initial explorations could include systems with microorganisms, where it has been very recently suggested that these complex cyclic assembly dynamics might occur (Song 2025). If our predictions hold, we envision that they would also advance our understanding of the relationship between multi-species assembly cycles and coalescence, a process in which entire microbial communities are replaced by others (Rillig et al. 2015; Diaz-Colunga et al. 2022).

At a macroscopic level, our findings explain the rarity of cycles in nature, at least for the empirical plant communities analyzed, based on two features commonly observed in ecological communities that are often coupled: intraspecific competition stronger than interspecific competition (Levine and HilleRisLambers 2009; Adler et al. 2018), and differences in species’ intrinsic growth rates (Godoy, Stouffer, et al. 2017; Ranjan et al. 2024). Seeing it from the opposite perspective, this coupling indicates that the presence of cycles and their frondosity (i.e., the number of nodes contained in the assembly graph) are constrained to ecological configurations governed by high interaction asymmetries and low variability in intrinsic growth rates. Our results, therefore, go beyond the classic dichotomy of absence/presence of cycles based on no universal best competitor. Assembly cycles are related to a more nuanced perspective of the complementarity of processes governing species population dynamics.

The term “cycle” has encompassed a broad range of perspectives and dynamics in the ecological literature. Many empirical approaches have documented cycles in the interaction matrix, also referred to as intransitivity, but these are not always linked to oscillatory dynamics in species abundances (Godoy, Stouffer, et al. 2017; Matías et al. 2018; Gallien, Zimmermann, et al. 2017). Others have studied such oscillatory dynamics but in tournaments (Laird et al. 2006; Soliveres, Maestre, et al. 2015; Allesina et al. 2011). The connection between cycles in the interaction matrix and oscillatory dynamics has been done through the study of heteroclinic and limit cycles (May et al. 1975; Hofbauer and So 1994; Gilpin 1975; Krupa and Melbourne 1995), but such studies have excluded assembly cycles occurring via successful invasions. Our study of cycles through the lens of assembly graphs should not be viewed as another independent approach. Instead, it groups all these previous cycles’ perspectives as different elements of an assembly graph. That is, it can contain internal oscillatory dynamics in a single sub-community as well as cyclic connections between sub-communities through invasions due to the lack of a universal best competitor. Therefore, we present here a general framework for cycles in ecology. This framework provides clear expectations of when cycles should be rare in nature and the associated consequences for the assembly and dynamics of ecological communities.

## Acknowledgments

The authors would like to express their gratitude for the financial support received. JAL and JRP acknowledge partial support from the Ministerio de Ciencia e Innovación, Spain, under Grant #PID2021-122991NB-C21. ROM acknowledges the support from the São Paulo Research Foundation (FAPESP) through Grants #2022/04886-2 and #2023/11798-5. VCS and OG acknowledge the financial support provided by the Spanish Ministry of Innovation, Science, and Universities and the European Social Fund through grants TASTE (#PID2021-127607OB-I00), and BIOTA (#EUR2023-143472). The authors thank the insightful comments of Carlos Gómez Ambrosi.

## Data accessibility statement

The data and code supporting the results have been archived at the repository: 10.5281/zenodo.14956574.

## Notes

### Competing Interest Statement

The authors have declared no competing interest.

### Summary of Updates

This version of the manuscript has been revised to update the following: improved flow, revisited structure, and added Supplementary information

## References

Adler, P. B. et al. (2018). “Competition and coexistence in plant communities: intraspecific competition is stronger than interspecific competition”. In: Ecol. Lett. 21.9, pp. 1319–1329.

Alcántara, J. M., M. Pulgar, and P. J. Rey (2017). “Dissecting the role of transitivity and intransitivity on coexistence in competing species networks”. In: J. Theor. Biol. 10, pp. 207–215.

Allesina, S. and J. M. Levine (2011). “A competitive network theory of species diversity”. In: Proc. Natl. Acad. Sci. U.S.A. 108.14, pp. 5638–5642.

Almaraz, P. et al. (2024). “Structural stability of invasion graphs for Lotka–Volterra systems”. In: J. Math. Biol. 88.6, p. 64.

Beninca, E. et al. (2015). “Species fluctuations sustained by a cyclic succession at the edge of chaos”. In: Proc. Natl. Acad. Sci. U.S.A. 112.20, pp. 6389–6394.

Bimler, M. D. et al. (2024). “Plant interaction networks reveal the limits of our understanding of diversity maintenance”. In: Ecol. Lett. 27.2, e14376.

Bortolan, M. et al. (2025). “A theoretical and computational study of heteroclinic cycles in Lotka–Volterra systems”. In: J. Math. Biol. 90.3, pp. 1–31.

Bunin, G. (2017). “Ecological communities with Lotka-Volterra dynamics”. In: Physical Review E 95.4, p. 042414.

Calleja-Solanas, V. et al. (2022). “Structured interactions as a stabilizing mechanism for competitive ecological communities”. In: Phys. Rev. E 106.6, p. 064307.

Chesson, P. (2000). “Mechanisms of maintenance of species diversity”. In: Annu. Rev. Ecol. Evol. Syst. 31.1, pp. 343–366.

Daniel, C. et al. (2024). “Fast–slow traits predict competition network structure and its response to resources and enemies”. In: Ecol. Lett. 27.4, e14425.

Diaz-Colunga, J. et al. (2022). “Top-down and bottom-up cohesiveness in microbial community coalescence”. In: Proceedings of the National Academy of Sciences 119.6, e2111261119.

Gallien, L., M. C. Cavaliere, et al. (2024). “Intransitive Stability Collapses Under the Influence of Dominant Competitors”. In: Am. Nat. 204.1, E000–E000.

Gallien, L., P. Landi, et al. (2018). “Emergence of weak-intransitive competition through adaptive diversification and eco-evolutionary feedbacks”. In: J. Ecol. 106.3, pp. 877–889.

Gallien, L., N. E. Zimmermann, et al. (2017). “The effects of intransitive competition on coexistence”. In: Ecol. Lett. 20.7, pp. 791–800.

Gilpin, M. E. (1975). “Limit cycles in competition communities”. In: Am. Nat. 109.965, pp. 51–60.

Godoy, O., F. Soler-Toscano, et al. (2024). “The assembly and dynamics of ecological communities in an ever-changing world”. In: Ecol. Monogr. 94.4, e1633.

Godoy, O., D. B. Stouffer, et al. (2017). “Intransitivity is infrequent and fails to promote annual plant coexistence without pairwise niche differences”. In: Ecology, pp. 1193–1200.

Hofbauer, J. and S. Schreiber (2022). “Permanence via invasion graphs: Incorporating community assembly into Modern Coexistence Theory”. In: J. Math. Biol. 85, p. 54.

Hofbauer, J. and J.-H. So (1994). “Multiple limit cycles for three dimensional Lotka-Volterra equations”. In: Appl. Math. Lett. 7.6, pp. 65–70.

Krupa, M. (1997). “Robust heteroclinic cycles”. In: J. Nonlinear Sci. 7, pp. 129–176.

Krupa, M. and I. Melbourne (1995). “Asymptotic stability of heteroclinic cycles in systems with symmetry”. In: Ergod. Theory Dyn. Syst. 15.1, pp. 121–147.

Laird, R. A. and B. S. Schamp (2006). “Competitive intransitivity promotes species coexistence”. In: Am. Nat. 168.2, pp. 182–193.

Levine, J. M., J. Bascompte, et al. (2017). “Beyond pairwise mechanisms of species coexistence in complex communities”. In: Nature 546.7656, pp. 56–64.

Levine, J. M. and J. HilleRisLambers (2009). “The importance of niches for the maintenance of species diversity”. In: Nature 461.7261, pp. 254–257.

MacArthur, R. (1969). “Species packing and what interspecies competition minimizes”. In: Proc. Natl. Acad. Sci. U.S.A. 64, pp. 1369–1375.

Matías, L. et al. (2018). “An experimental extreme drought reduces the likelihood of species to coexist despite increasing intransitivity in competitive networks”. In: J. Ecol. 106.3, pp. 826–837.

May, R. and W. Leonard (Sept. 1975). “Nonlinear Aspects of Competition Between Three Species”. In: SIAM J. Appl. Math. 29, p. 243.

McCann, K., A. Hastings, and G. R. Huxel (1998). “Weak trophic interactions and the balance of nature”. In: Nature 395.6704, pp. 794–798.

Meyer, C. D. (2023). Matrix analysis and applied linear algebra. SIAM.

Neutel, A.-M., J. A. P. Heesterbeek, and P. C. de Ruiter (2002). “Stability in Real Food Webs: Weak Links in Long Loops”. In: Science 296.5570, pp. 1120–1123.

Pajares-Murgó, M. et al. (2024). “Intransitivity in plant–soil feedbacks is rare but is associated with multispecies coexistence”. In: Ecol. Lett. 27.3, e14408.

Ranjan, R., T. Koffel, and C. A. Klausmeier (2024). “The three-species problem: Incorporating competitive asymmetry and intransitivity in modern coexistence theory”. In: Ecol. Lett. 27.4, e14426.

Rillig, M. C. et al. (2015). “Interchange of entire communities: microbial community coalescence”. In: Trends in Ecology & Evolution 30.8, pp. 470–476.

Rojas-Echenique, J. and S. Allesina (2011). “Interaction rules affect species coexistence in intransitive networks”. In: Ecology 92.5, pp. 1174–1180.

Saiz, H. et al. (2019). “Intransitivity increases plant functional diversity by limiting dominance in drylands worldwide”. In: J. Ecol. 107.1, pp. 240–252.

Serván, C. and S. Allesina (2021). “Tractable models of ecological assembly”. In: Ecol. Lett. 24, pp. 1029–1037.

Soliveres, S. and E. Allan (2018). “Everything you always wanted to know about intransitive competition but were afraid to ask”. In: J. Ecol. 106.3, pp. 807–814.

Soliveres, S., F. T. Maestre, et al. (2015). “Intransitive competition is widespread in plant communities and maintains their species richness”. In: Ecol. Lett. 18.8, pp. 790–798.

Song, C. (2025). “Assembly Graph as the Rosetta Stone of Ecological Assembly”. In: Environ. Microbiol. 27.1. e70030 EMI-2024-0920.R2, e70030.

Takeuchi, Y. (1996). Global Dynamical Properties of Lotka–Volterra Systems. Singapore: World Scientific Publishing Co. Pte. Ltd.

Tilman, D. (1982). Resource competition and community structure. 17. Princeton university press.

Tylianakis, J. M. et al. (2010). “Conservation of species interaction networks”. In: Biol. Conserv. 143.10, pp. 2270–2279.

Ulrich, W. et al. (2018). “Functional traits and environmental characteristics drive the degree of competitive intransitivity in European saltmarsh plant communities”. In: J. Ecol. 106.3, pp. 865–876.

Vandermeer, J. (2011). “Intransitive loops in ecosystem models: from stable foci to heteroclinic cycles”. In: Ecol. Complex. 8.1, pp. 92–97.

(2013). “Forcing by rare species and intransitive loops creates distinct bouts of extinction events conditioned by spatial pattern in competition communities”. In: Theoretical ecology 6, pp. 395–404.

(2025). “Adding species to an intransitive triplet: competition, hierarchy, and recoupling”. In: J. Theor. Biol. 18.1, p. 8.

Vandermeer, J. and I. Perfecto (2023). “Intransitivity as a dynamic assembly engine of competitive communities”. In: Proc. Natl. Acad. Sci. U.S.A. 120.15, e2217372120.

Wootton, J. T. (1994). “The nature and consequences of indirect effects in ecological communities”. In: Annu. Rev. Ecol. Evol. Syst., pp. 443–466.

Zeeman, M. L. (1993). “Hopf bifurcations in competitive three-dimensional Lotka–Volterra systems”. In: Dyn. Stab. Syst. 8.3, pp. 189–216.

