## Supplementary S1, S2 for "A general framework for cycles in ecology"

#### Appendix S1: Algorithm for generating Volterra-Lyapunov stable matrices

#### Appendix S2: Algorithm for generating Volterra-Lyapunov stable matrices containing *à la carte* cyclic behavior

### S1. Algorithm for generating Volterra-Lyapunov stable matrices

Almaraz et al. 2024 has recently demonstrated that for Lotka–Volterra systems with Volterra–Lyapunov (VL) stable interaction matrices, the assembly graph is equivalent to the graph that represents the connections between the system equilibria. Building on this parallelism, our study focuses on systems with VL stable matrices, as they provide a clear mathematical framework for analyzing cycles. Since our primary goal is to explore the conditions under which cycles emerge, we needed an efficient way to generate a large number of VL stable matrices. To address this, we use a characterization of VL stability that enables either the random generation of VL stable matrices or their construction according to specific research objectives (Bortolan et al. 2025). In particular, we found that a *Volterra-Lyapunov* stable matrix $A$ can be written in the form:

$$A = H\,(S + J) \tag{1}$$

where $S$ is a symmetric and stable matrix (i.e. all its eigenvalues are strictly negative), $H = diag(h_i)$ with all $h_i > 0$, and $J$ is any anti-symmetric matrix. To generate VL stable matrices, we first draw a random positive diagonal matrix $H$. Next, we create a random anti-symmetric matrix $J$ by starting with a random matrix $K$ and taking $J = K - K^T$. Finally, we construct a random symmetric stable matrix $S$ by generating another random matrix $F$ and calculating $S = F + F^T - \alpha * \mathbb{I}$, where $\alpha \geq 0$ is chosen to be sufficiently large so that all eigenvalues of $S$ are strictly negative, and $\mathbb{I}$ is the identity matrix. This procedure guarantees that we can obtain a wide range of matrices $A$ with varying asymmetry ratios, and all of them meet the mathematical criteria for VL stability.

### S2. Algorithm for generating Volterra-Lyapunov stable matrices containing *à la carte* cyclic behavior

For illustrative purposes of our main findings, it is useful to create an interaction matrix with a desired cyclic behavior (see Figs. ??, ?? and ??). Here, we introduce an algorithm capable of doing this by extending the classic three-species competitive community of May and Leonard (May et al. 1975) into five dimensions. This extension generates an assembly graph with a cycle of five single-species nodes known as the rock-paper-scissors-lizard-Spock (RPSLS) game. To create this assembly graph, we use the characterization a Volterra-Lyapunov stable matrix given in Eq. (1), setting $H$ equal to the identity matrix $\mathbb{I}$. We can then write a matrix as $A = A_S(d) + A_J$, where $d < 0$ and the symmetric and anti-symmetric components are given by:

$$A_S(d) = \begin{pmatrix} d & 0 & 0 & 0 & 0 \\ 0 & d & 0 & 0 & 0 \\ 0 & 0 & d & 0 & 0 \\ 0 & 0 & 0 & d & 0 \\ 0 & 0 & 0 & 0 & d \end{pmatrix} = d\mathbb{I},$$

$$A_J = \begin{pmatrix} 0 & -1 & 1 & -1 & 1 \\ 1 & 0 & -1 & 1 & -1 \\ -1 & 1 & 0 & -1 & 1 \\ 1 & -1 & 1 & 0 & -1 \\ -1 & 1 & -1 & 1 & 0 \end{pmatrix}$$

The anti-symmetric matrix $A_J$ represents the positive and negative interactions in an RPSLS game and intrinsic growth rates were all chosen to be 1. The increase of the absolute value of $d$ makes the system more symmetric, as the species interact more within themselves than with the others. Since we have predicted that a diagonally dominant matrix inhibits cycles, which occur when $d < -4$, we chose $|d|$ to be small so that the symmetry of the matrix does not inhibit the cyclic behavior. In this example, cycles exist in the assembly graph when $d \in (-1, 0)$, but they disappear when $d = -1$.

Extending the ideas of this previous example, we can create interesting cyclic behavior for any dimension. The core of the approach is to sum a diagonal stable matrix like $A_S(d)$ with an anti-symmetric matrix $A_J$ containing the cyclic interactions between the species. When the diagonal terms of $A_S(d)$ are small in absolute values, we have a small symmetric part and the cycles emerge in the assembly graph. Finally, for species-varying cycles (the third category in Figure **??** C), suppose we want a cycle where species 3 beats species 1 and 4, then species 4 beats 3, and species 1 and 2 beat 4. This behavior is present in the dynamics of the system with the matrix:

$$A_{double} = \begin{pmatrix} d & 0 & -1 & 0.5 \\ 0 & d & -1 & 0.5 \\ 1 & 1 & d & -1 \\ -0.5 & -0.5 & 1 & d \end{pmatrix}$$

The matrix $A_{double}$ is Volterra-Lyapunov stable, and its anti-symmetric component represents the cyclic dynamics we want to create (cycles between one-species and two-species subcommunities). When $d \in (-1, 0)$, we have the cycle $\{1, 2\} \to \{3\} \to \{4\} \to \{1, 2\}$ in the assembly graph of the system. When $d = -1$, the cycle breaks, and when $d < -1$, the assembly graph of the system becomes identical to the assembly graph of the decoupled symmetric system with matrix $d \cdot \mathbb{I}$.

### References

Almaraz, P. et al. (2024). "Structural stability of invasion graphs for Lotka–Volterra systems". In: *J. Math. Biol.* 88.6, p. 64.

Bortolan, M. et al. (2025). "A theoretical and computational study of heteroclinic cycles in Lotka–Volterra systems". In: *J. Math. Biol.* 90.3, pp. 1–31.

May, R. and W. Leonard (Sept. 1975). "Nonlinear Aspects of Competition Between Three Species". In: *SIAM J. Appl. Math.* 29, p. 243.
